# SCIE: Information Extraction for Spinal Cord Injury Preclinical Experiments – A Webservice and Open Source Toolkit

**DOI:** 10.1101/013458

**Authors:** Andreas Stöckel, Benjamin Paassen, Raphael Dickfelder, Jan Philip Göpfert, Nicole Brazda, Hans Werner Müller, Philipp Cimiano, Matthias Hartung, Roman Klinger

**Affiliations:** Semantic Computing, CITEC, Bielefeld University; Institute for Natural Language Processing (IMS), University of Stuttgart; Neurology, HHU Düsseldorf and Center for Neuronal Regeneration Düsseldorf

## Abstract

Translational neuroscience in the field of spinal cord injuries (SCI) faces a strong disproportion between immense preclinical research efforts and a lack of therapeutic approaches successful in human patients: Currently, preclinical research on SCI yields more than 3,000 new publications per year (8,000 when including the whole central nervous system, growing at an exponential rate), whereas none of the resulting therapeutic concepts has led to functional recovery of neural tissue in humans. Improving clinical researchers’ information access therefore carries the potential to support more effective selection of promising therapy candidates from preclinical studies. Thus, automated information extraction from scientific publications contributes to enabling meta studies and therapy grading by aggregating relevant information from the entire body of previous work on SCI.

We present SCIE, an automated information extraction pipeline capable of detecting relevant information in SCI publications based on ontological entity and probabilistic relation detection. The input are plain text or PDF documents. As output, the user choses between an online visualization or a machine-readable format. Compared to human gold standard annotations, our system achieves an average extraction performance of 76 % precision and 52 % recall (F_1_-measure 0.59).

An instance of the webservice is available at http://scie.sc.cit-ec.uni-bielefeld.de/. SCIE is free software licensed under the AGPL and can be downloaded for local installation at http://opensource.cit-ec.de/projects/scie/.

## 1 Introduction

Injury to the central nervous system of adult mammals typically results in lasting deficits, like permanent motor and sensory impairments, due to a lack of profound neural regeneration. Specifically, patients who have sustained spinal cord injuries (SCI) usually remain partially paralyzed for the remainder of their lives. Preclinical research in the field of central nervous system trauma advances at a fast pace, currently yielding over 8,000 new publications per year, at an exponentially growing rate, with a total amount of approximately 160,000 PubMed-listed papers today.^1^

However, translational neuroscience faces a strong contrast between the amount of preclinical knowledge on SCI published in scientific articles, on the one hand, and the lack of successful clinical trials in SCI therapy, on the other: So far, no therapeutic approach has led to functional recovery in human patients (Filli and Schwab, 2012). As the vast amount of published information by far exceeds the capacity of individual scientists to read and absorb the relevant knowledge (Lok, 2010), selection of promising therapeutic interventions for clinical trials is notoriously based on incomplete information (Steward et al., 2012). Thus, automatic information extraction methods are needed to gather comprehensive actionable knowledge in structured form from scientific articles that describe outcomes of preclinical experiments in the SCI domain.

We present SCIE, an implementation of an information extraction system for acquisition of structured knowledge about experimental SCI therapies from research papers. It uses an ontology-based named entity recognition (NER) and a machine learning-based slot-filling module for the detection of complex relationships. The webservice supports interactive visualization and exporting to a JSON format. In addition, a command line application is available for integration of SCIE into existing UIMA workflows.^2^

## 2 Methods

Scientific articles in the SCI domain typically contain four important *relations*: a group of laboratory ANIMALs, an INJURY inflicted on the animals, a TREATMENT and the observed RESULT (Brazda et al., 2013). We phrase the relation extraction as slot filling of templates: *e. g.* the Animal template provides slots for Organism, Laboratory ANIMAL, Weight, Age, Sex and Number (*cf*. Fig. 2).

An illustration of the entire SCIE workflow is shown in Fig. 1. In case of PDF input, the document is transformed into plain text (based on Apache PDFBox^3^). *Named entities* are extracted based on regular expressions to detect entities such as quantities and units and on ontologies with approximative string matching comprising token lookup, candidate generation and filtering as well as an ontological reduction (Paassen et al., 2014). Animal names are extracted according to the vertebrates subgraph within the NCBI Taxonomy^4^ and drug names as in MeSH^5^. Domain-specific knowledge (*e. g.*, injury methods) is covered by manually tailored ontologies.

**Figure 1:**
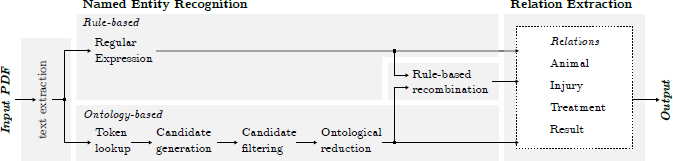
The workflow of SCIE consists of text extraction, named entity recognition and relation detection.

**Figure 2:**
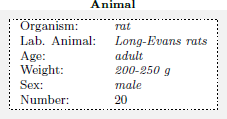
An example of the ANIMAL relation with slots and entities.

**Figure 3:**
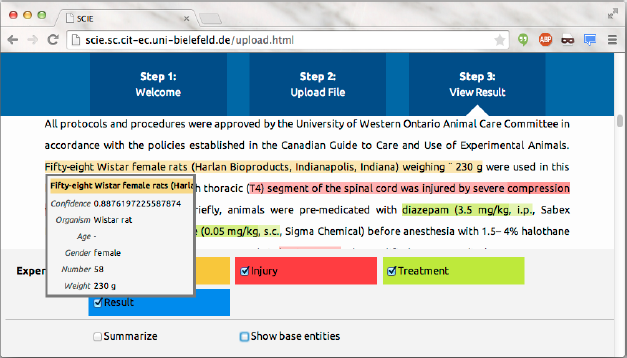
Screenshot of the outcome of our system processing a text snippet from Gris et al. (2005).

*Relation extraction* aggregates the previously identified entities in a slot filling setting into ANIMAL, INJURY, TREATMENT and RESULT templates. All possible candidates of entity combinations are ranked using a learning-to-rank approach (Liu, 2009) based on logistic regression which assigns each relation candidate a confidence score. The models have been trained on a manually annotated corpus of 18 full papers. After ranking all relation candidates, the six best-ranked results for each relation are returned to the user.

## 3 Software and Interfaces

SCIE is composed of independent and reusable modules (PDF extraction, NER, Relation Extraction). The SCIE core application implements the rule-based NER and provides a command line interface for batch-processing and UIMA XCAS output. The webservice is based on an embedded Jetty HTTP Server^6^ and provides a user interface for uploading PDF and plain text documents, running an analysis and viewing the resulting annotations and an HTTP API for batch processing of documents and downloading of results in a JSON format^7^. The webservice module itself does not depend on UIMA or the SCIE core application, yet a corresponding binding module is provided.

## 4 Results and Discussion

The *F*_1_ measure for retrieving entities participating in one of the four relations is 0.59 (0.76 precision, 0.52 recall). For animal, it is 0.82 (0.96/0.74), for Injury 0.72 (0.71/0.73), for Treatment 0.47 (0.67/0.37) and for Results 0.38 (0.68/0.27) based on a set of 32 manually annotated articles. The offsets and frequencies of the annotations of entities have not been taken into account. Processing all 32 papers took 5 minutes and 45 seconds on an AMD Phenom quad-core clocked at 3 GHz with 4 GB RAM (10.8 seconds/paper).

SCIE provides an easy-to-use web interface (*cf.* Suppl. Material) that can assist medical researchers in finding the relevant sections in research papers in the SCI domain. Applying the API workflow^8^ to automatically populate a database will enable access to a comprehensive pool of SCI domain knowledge far beyond individual publications, thus paving the way towards complex meta studies and empirical therapy grading (Kwon et al., 2011).

Future work includes the enrichment of the training data to further improve the performance of the automatic information extraction. In addition, we will extend our pipeline towards an information retrieval system and ultimately populate a database.

## Acknowledgement

Roman Klinger has been partially funded by the “It’s OWL” project (“Intelligent Technical Systems Ostwestfalen-Lippe”), a leading-edge technology and research cluster funded by German Ministry of Education and Research (BMBF).

## Footnotes

1 As in this query (http://tinyurl.com/sciquery) to http://www.ncbi.nlm.nih.gov/pubmed, as of November 2014.

2 https://uima.apache.org/

3 Apache PDFBox – A Java PDF Library (http://pdfbox.apache.org/)

4 NCBI Taxonomy (http://www.ncbi.nlm.nih.gov/taxonomy)

5 NCBI Medical Subject Headings, (http://www.ncbi.nlm.nih.gov/mesh)

6 Jetty – Servlet Engine and Http Server (http://www.eclipse.org/jetty/)

7 JSON (http://www.json.org/)

8 http://scie.sc.cit-ec.uni-bielefeld.de/api.html

